# One-week exposure to South Indian Classical music clip having incremental variation in tempo and octave promotes better anxiety reduction among medical students – an EEG based study

**DOI:** 10.1101/656777

**Authors:** Sushma Sharma, Arun Sasidharan, Vrinda Marigowda, Mohini Vijay, Sumit Sharma, Chetan Satyajit Mukundan, Lakshmi Pandit

## Abstract

Several scientific studies using Western classical music and some using Indian classical music have reported benefits of listening to musical pieces of specific ‘genre’ or ’Raga’, in terms of stress reduction and mental well-being. Within the realm of a Raga, presentation of musical pieces varies in terms of low-level musical components (like tempo, octave, timbre, etc.), and yet there is hardly any research on their effect. A commonly preferred musical pattern in Carnatic classical music is to have incremental modulations in tempo and octave (‘*Ragam-Tanam-Pallavi*’), and we wanted to examine whether this could have better anxiolytic effect than music without such modulations.

Accordingly, in the current study, we exposed 21 male undergraduate medical students to a custom recorded South Indian classical music clip for 1 week (8 minutes clip; Raaga ‘*Kaapi*’; only two instruments – ‘Violin’ and ‘Mridangam’; listened thrice daily for 6 days). One set of the participants (Varying Music; n=11) listened to a version that had the incremental variations, whereas the other set (Stable Music; n=10) listened to a version without such variations. On all 6 days, one of the music listening sessions was conducted in the lab while collecting electroencephalography (EEG; 32 channels) and electrocardiography (ECG; 1 channel) data. Psychological assessment for anxiety (State-Trait Anxiety Inventory - STAI and Beck Anxiety Inventory - BAI) was conducted before (day 1) and after (day 6) the intervention. Physiological parameters studied included power spectrum across the scalp in delta, alpha, beta, theta and gamma bands from EEG and heart rate variability (HRV) from ECG, during the baseline recordings of day 1 and day 6 of intervention.

Our results show that participants when exposed to varying music showed significant reduction in anxiety, in contrast to stable music or silence intervention. A global examination of power spectral changes showed a stark contrast between stable and varying music intervention in comparison to silence - former showing greater increase in higher frequencies whereas latter showing prominent decrease especially in lower frequencies, both in bilateral temporo-parieto-occipital regions. A more detailed spectral analysis in frontal region revealed that both music intervention showed greater left-dominant alpha/beta asymmetry (i.e., greater right brain activation) and decrease in overall midline power (i.e., lower default mode network or DMN activity), when compared to silence intervention. Interestingly, stable music resulted in more left asymmetry, whereas, varying music showed more midline power reduction. Both music intervention also didn’t show the reduction in HRV parameters that was associated with silence intervention.

We speculate that, the enhancement in ‘mind calming effect’ of *Kaapi raaga* when presented with incremental variations, could be brought about by a balanced switching between a heightened mind wandering state with ‘attention to self’ during the lower-slower portions and a reduced mind wandering state with ‘attention to music’ during the higher-faster portions of the music. Such a ‘dynamic mind wandering’ exercise would allow training one’s creative thinking as well as sustained attention, during the respective high and low mind wandering states - both helping prevent ruminating thoughts. Therefore, musical properties such as tempo and octave have non-trivial influence on the various neurological and psychological mechanisms underlying stress management. Considering the impact of this finding in selection of music clips for music therapy, further studies with larger sample size is warranted.

## 2. INTRODUCTION

Stress has now become an ingrained part of our vocabulary and daily existence. Hans Seyle described Stress as a biopsychosocial model that refers to the consequence of failure of an organism to respond adequately to mental, emotional and physical demands, whether actual or imagined (Selye, 1982). A stressful situation is one in which demands of the situation threaten to exceed the resources of the individual (Lazarus & Folkman, 1984). Students are subjected to different kinds of stressors such as pressure of the academics with an obligation to succeed, an uncertain future and difficulties of integrating into the system (Fish & Nies, 1996). Medical education itself is reported to be extremely demanding and stressful (Firth-Cozens, 1999; Lee & Graham, 2001) and medical curriculum can contribute to the development of depression and anxiety related problems in the medical students which may have possible negative academic consequences. Indian studies have reported that overall prevalence of depression, anxiety and stress among medical students was found to be 71.23% (G. Kumar, Jain, & Hegde, 2012), 66.9% and 53% respectively (Iqbal, Gupta, & Venkatarao, 2015). In the Indian scenario, the important determinants of stress among medical students were found to be number of years of studying the course, vastness of academic curriculum, fear of poor performance in examination, lack of recreation, high workload and family problems (Anuradha, Dutta, Raja, Sivaprakasam, & Patil, 2017). Although some stress is expected in college and it can be a motivation to study and learn, too much stress can deter learning and have deleterious effects on health and cognitive functioning of students (V. Kumar, Talwar, & Raut, 2014). Exposure to intense and chronic stressors has long lasting neurobiological effects and puts one at increased risk of anxiety and mood disorders, aggressive dyscontrol problems, hypo immunity, medical morbidity, structural changes in the central nervous system and early deaths (Shaw, 2003).

Studies have routinely documented that states of enduring low mood are often associated with mind wandering (Smallwood, O’Connor, Sudbery, & Obonsawin, 2007). Mind wandering can be defined as the conscious processing of information that is unrelated to immediate sensory input and to the task currently being performed (Smallwood, Mrazek, & Schooler, 2011). Mind wandering generally impairs learning, increases the amount of off-topic thought and results in absent minded errors (Smallwood, Fitzgerald, Miles, & Phillips, 2009). This would imply that students who mind wander frequently are likely to underperform in academics (Smallwood, Fishman, & Schooler, 2007). Given the fact that the medical community is more vulnerable to stress and anxiety related problems it may be argued that mind wandering may be relatively frequent within the medical community.

Mindfulness is a state of mind in which the individual is aware of his own mental processes, listen more attentively and thereby act with principle and compassion (Epstein, 1999).

Mindfulness has the potential to prevent compassion fatigue and burnout because the doctor who is self-aware is more likely to engage in self-care activities and to manage stress better.

Therefore, it is essential to propose and develop new strategies that are acceptable and easily adaptable to help individuals become more mindful and promote overall psychological wellbeing. This could ameliorate the cost of the problems to the medical community and society at large.

Lifestyle changes that decrease stress and anxiety are thought to be highly effective (Epstein, 1999) and music therapy may be one among them (Dileo & Bradt, 2009). The vast body of existing literature reveals that listening to music has the inherent ability to relieve stress and promote overall well-being of an individual (Cepeda, Carr, Lau, & Alvarez, 2006; Dileo & Bradt, 2009; Epstein, 1999; Harvey, 1987; Iwanaga, Ikeda, & Iwaki, 1996; Nilsson, Unosson, & Rawal, 2005; Phipps, Carroll, & Tsiantoulas, 2010). Music therapy enhances attention and mood, particularly the mood state of fatigue and thus can be considered as a mindfulness based interventional tool (Lesiuk, 2015). Listening to classical music has higher effects on stress reduction compared to non-classical music such as Hard rock and Heavy metal music (H. Hasegawa, Uozumi, & Ono, 2003). Moreover, listening to instrumental music has shown to induce greater sense of relaxation than music combined with lyrics (Burns et al., 2002).

With the advent of modern neuroimaging techniques, there is an increasing interest in uncovering the neural structures associated with music perception. EEG is a time-tested method used for studying the transient dynamics of human brain’s large-scale neuronal circuits. The alpha waves (8-12Hz) seem to be accompanied by characteristic tranquil or calm states of mind, beta waves (12-30Hz) are considered as sensory motor rhythm and gamma waves (30-80 Hz) seem to be associated with a wide range of cognitive functions. A study on Indian classical music has shown that listening to raga ‘Desi Thodi’ has led to a significant increase in the alpha EEG frequency and a significant decrease in scores on depression, state and trait anxiety (Gupta & Gupta, 2005).

Besides the high-level grouping of music into genre, it can also be described in terms of components like tempo, mode, octave, rhythm, timbre, harmony, melody and loudness (Amezcua, Guevara, & Ramos-Loyo, 2005). To fully understand the psychological effects of music, it is vital to dissect and isolate the individual components of music (Kellaris & Kent, 1991; Oakes, 2003). Of all the music properties, tempo is of greatest importance in carrying the expressiveness in music (Hevner, 1937). Tempo communicates the pace of the music; it is measured as beats per minute and is regarded as the most important feature in determining the mood effects of music (Rigg, 1964). Changes in musical tempo evokes movement-related brain activity (Daly et al., 2014), produces significant changes in EEG alpha power (Ma, Lai, Yuan, Wu, & Yao, 2012), stimulates arousal level and varies vigilance of an individual (C. Hasegawa & Oguri, 2006) and can also influence mind wandering depending on the type of music (sad or happy) (Taruffi, Pehrs, Skouras, & Koelsch, 2017). Moreover, happy and joy musical excerpts exhibit greater left frontal EEG activity whereas fear and sad musical excerpts exhibited relative right frontal EEG activity (Schmidt & Trainor, 2001). Musical octave is another important musical property, which can be defined as the interval between one musical pitch and another with half or double its frequency. Very limited information is available regarding the perception of musical octave and its effects on stress response. Finally, majority of the previous work has only focused on examining the overall positive effects of music on stress response and a very little attempt has been made in the past to understand the importance of individual music properties in the management of chronic stress.

In the light of the above lacunae in existing research, we set out to examine the effects of varying both musical tempo and octave, on physiological and psychological parameters related to stress management. We analysed physiological changes using the EEG spectral power and heart rate variability, and subjective perception of anxiety by using psychological questionnaires. *Ragam-Tanam-Pallavi* (RTP) is one of the most commonly performed musical pieces in a South Indian (Carnatic) Classical music concert. The RTP begins by a section in free rhythm to create the mood of raga (‘*Raga alapana*’), moves into a more rhythmic pattern characterized by a gradual increase in tempo (‘*Tanam*’), and progresses to an overall melodic ascent that ends in rapid passages. The first stage mostly begins in a slow tempo and melodic features in the lower register and the final stage ends with high tempo in upper register (Henry, 2002). In accordance with this concept of RTP, a music clip was developed that had a crescendo pattern of variation in tempo and octave. We called this music with incremental modulations in tempo and octave as Varying music (VM) and the music with no such variations as Stable music (SM) and we chose Violin for melody and Mridangam for percussion in both the music types.

In this study, we hypothesized that participants listening to VM would show a different physiological and psychological response when compared to participants listening to SM or a non-music control intervention (silence).

## 3. MATERIALS AND METHODS

### 3.1. Subjects

A total of 25 healthy right-handed (based on self-report) male undergraduate medical students aged 18-24 years were recruited as subjects.

We included only male subjects in this study as hormonal variations occurring within menstrual phases are known to affect a female’s response on various psychophysiological measures.

None of the participants had received any formal musical training, had a history of mental or neurological problems, hearing impairments, prior exposure to the tested music or any form of music therapy. All participants signed an informed consent form and study protocol that was approved by the institute human ethics committee.

### 3.2. Music stimulus

In the present study Indian classical music played by Violin and Mridangam in the raga ‘*Kaapi*’ was chosen. The music clip was custom recorded for this study by two experts in Indian carnatic music, who were blind to this study. Two different music pieces played by the above-mentioned instruments were selected. One had an incremental variation in the tempo and the octave (VM) and the other had a more stable tempo and octave (SM). The duration of both the music pieces was about eight minutes.

The music piece for VM: First forty seconds (0-40s) had the violin played in lower octave and Mridangam played with one beat per second. The next forty seconds (40-80s) had the Violin played in middle octave and Mridangam played with two beats per second. The next forty seconds (80-120s) had the Violin played in upper octave and Mridangam played with four beats per second. This 120 second pattern was repeated 3 more times to make up the 8 minutes music clip. Thus, the music clip had 4 incremental sequences.

The music piece for SM: Throughout eight minutes of music, the tempo and octave were kept constant without much variations; i.e., Mridangam was played with one beat per second and violin was played in only middle octave.

### 3.3. Experimental Design

A cross-over study design was followed. The subjects were randomly assigned to have one of the two different music interventions (VM or SM) and a control intervention (silence), each for six days; music and control sessions were counterbalanced and spaced apart by at least 1 week. So, each subject in the experimental group served as his own control for the music vs silence comparison. 4 participants could not complete the whole study due to personal/technical reasons and are not included in further analysis. Thus, we had 11 participants for VM group and 10 for SM group. The counterbalancing of music therapy and silence exposure was nearly equal in both the groups (5 in both music groups had music first and silence later, and the rest had silence first and music later).

During the week of music intervention, the participants listened to the assigned music clip thrice daily (morning, noon and evening) for 6 days. Many participants did not complete the intervention within 1 week. Considering practicality, the intervention was considered complete if there was no more than 2 days gap between the sessions and all the six intervention days were completed within 12 days of start of the intervention. Every day, for the evening music sessions (around 3PM-7PM), subjects were required to visit the laboratory and listen to music along with EEG recording. The participants sat in eyes-closed relaxed state throughout the session, in a dim lit and sound-attenuated experimental room; music was presented through speakers (at 55-65 dB based on personal preference). Music preference of the subjects was not considered in the present study. The subjects during the silence period followed the same procedure, except that no music was played.

EEG data was recorded using a saline-based 32-channel EEG electrode cap (RapidCap) and EEG-ERP system (BESS FW-32, Axxonet System Technologies Pvt Ltd., Bengaluru) following the 10-10 electrode position system. The data was digitized at a sampling frequency of 1024 Hz. After the music session, the subjects were asked to rate the music as well as their sleepiness during the session on a scale of 10. The subjects were asked to fill the psychological questionnaires on day 1 before the music session and day 6 after completing the music session.

### 3.4. Questionnaire-based Psychological Assessment

All participants were required to fill two questionnaires on the first day and last day of intervention (in both music as well as silence conditions). The State-Trait Anxiety Inventory (STAI) (Spielberger, Gorsuch, Lushene, Vagg, & Jacobs, 1983) and Beck Anxiety Inventory (BAI) (Beck, Epstein, Brown, & Steer, 1988) were chosen to measure anxiety. These tests are quick and easy to administer, known to have good reliability and validity, and can together have a broad coverage of general symptoms of anxiety (Julian, 2011).

### 3.5. EEG Paradigm

The experimental paradigm consisted of 3 runs of EEG recordings –(1) Pre-intervention recording which is period of silence for 3 minutes, (2) Intervention recording during which music presented for 8 minutes and (3) post-intervention recording which is again a period of silence for 3 minutes (Fig. 1).

**Figure 1:**
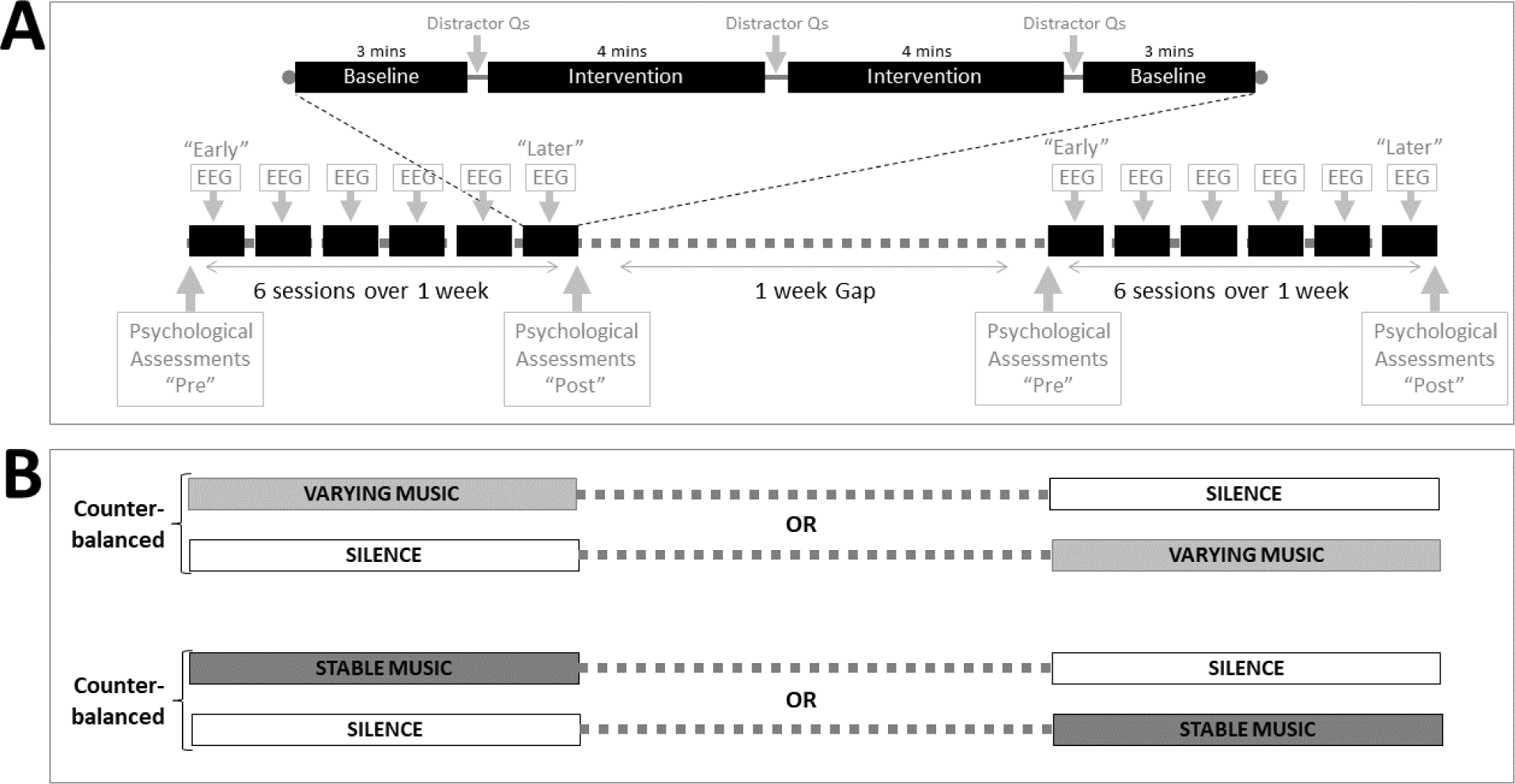
Study Design showing (A) detailed timeline of the sessions, and (B) counterbalancing of the cross-over design.

Simple arithmetic questions (a set of three questions; e.g. “Is two plus five equals to eight?”) were presented during the EEG session, and the subject was asked to press a keypad button for “yes” or “no”. This was included to prevent the subject from sleeping off and ensure engagement of the subject. First set of questions were asked right before the beginning of the music, the second set of questions were asked in the middle of music listening (i.e. after 4 mins of start of music clip) and the third set of questions were asked after the completion of music.

### 3.6. EEG Analysis

EEG data was first subjected to pre-processing using EEGLAB v13 toolbox (Delorme & Makeig, 2004) implemented in Octave (https://www.gnu.org/software/octave/). EEG drift artefacts removed using a 0.25-0.75Hz transition band high-pass filter, bad channels were rejected using an automated correlation based routine, and high amplitude stereotypical (e.g., eye blinks) and non-stereotypical (e.g., movement) artefacts removed using an artefact subspace reconstruction (ASR) algorithm (Mullen et al., 2013); all implemented using a plugin in EEGLAB. ASR relies on a sliding window principal component analysis (PCA) to statistically compare with a reference clean portion of the EEG data and then linearly reconstruct deviant portions of the data (‘artefact subspaces’) based on the correlation structure observed in the reference clean data (Mullen et al., 2013). We used a threshold of 10 standard deviations for ASR. Deleted bad channels were interpolated using spherical spline interpolation and re-referenced to all channel average.

EEG data from two time points, Day 1 (“Early”) and Day 6 (“Later”) (Fig 1), were used for power spectral analysis. In subjects whose Day 1 data was of poor quality, Day 2 was used, and similarly Day 5 was used instead of a poor-quality Day 6 data; 1 subject from each group had a bad day 1 or day 6 data. 31 channel artefact-free EEG data were then imported to Brainstorm (Tadel, Baillet, Mosher, Pantazis, & Leahy, 2011), an open-source EEG analysis software (http://neuroimage.usc.edu/brainstorm), as 10s non-overlapping epochs of each condition.

Overall 20 epochs (∼3 mins) of pre-music rest and during music/silence were selected from each subject, from the two time points. Each 10s epoch was subjected to power spectral density estimation using p-welch algorithm (2s hamming window; 50% overlap; 0,5Hz resolution).

The power spectral density values were then subjected to hierarchical general linear model analysis using the LIMO toolbox (Pernet, Chauveau, Gaspar, & Rousselet, 2011). Briefly, the data of each subject from each timepoint was subjected to mass univariate analysis with Rest and Music/Silence as categorical variables. This first-level analysis gave ‘beta’ values with reduced inter subject and inter-session variability, and was used for second-level (between-group) analysis. We compared the spectral changes between the two timepoints using robust paired t-test, for each group, as implemented in the LIMO toolbox. Cluster statistics (threshold-free cluster enhancement) (Smith & Nichols, 2009) was used to correct for multiple comparison effect across multiple channels and frequency.

Besides the group-level analysis as mentioned above, as subject-level analysis was done for frontal asymmetry and midline frontal power. To calculate frontal asymmetry, firstly, instantaneous amplitude envelope was extracted for each channel and frequency band across each epoch using hilbert transform approach. Those from right frontal (Fp2, F4, F8, FC4, FT8) and left frontal (Fp1, F3, F7, FC3, FT7) electrodes were separately averaged, log transformed and then subtracted to get asymmetry index. A cumulative sum of this difference was taken across each 10s epoch, whose slope was taken as the final measure of frontal asymmetry. This approach was better in capturing the overall trend of hemispheric asymmetry despite the numerous transitions between right and left dominance that is common over longer epochs. The subject-level change in frontal asymmetry values between early and later time points (separately for each intervention) was computed and their significance was determined using bootstrap statistics (1000 resamples; 95% confidence intervals). The significant increase and decrease in asymmetry were coded as ‘+1’ and ‘-1’ respectively, and all non-significant values as ‘0’, for further computation. The sum of these coded values for alpha and beta bands was used to make subject-level inference of which intervention showed greater frontal asymmetry.

The midline frontal power was also computed from the amplitude envelope, but from all frequency bands, and coded for subject-level inference as mentioned for frontal asymmetry.

### 3.7. Heart rate variability (HRV) Analysis

A single ECG channel (positioned on bilateral clavicles; lead-II) was used for HRV analysis. An automated algorithm was used for extracting RR intervals (Pan & Tompkins, 1985), which is available as a MATLAB toolbox (Sedghamiz, 2014). Standard and novel HRV parameters were computed using another MATLAB toolbox (HRVTool; www.MarcusVollmer.github.io/HRV). A frequency domain parameter (LF/HF ratio) was chosen as the standard measure and a new robust geometric of HRV based on relative RR intervals (rrHRV) was also computed (Vollmer, 2015).

### 3.8. Other Statistical Analysis

Each of the anxiety scores were subjected to 2-way ANOVA with group and sessions (or intervention type) as factors, followed by post-hoc t-tests with Tukey’s p-value correction.

## 4. Results

### 4.1. Anxiety score differences

When comparing anxiety scores between the first and last days of the 1-week intervention, significant group and session interaction was seen (F_3,37_=9.13, p=0.005). On post-hoc analysis, only VM group showed significant reduction in anxiety scores following music intervention when compared to silence intervention in all the three measures (STAI-state: t=-5.57, p<0.001; STAI-trait: t=-5.67, p<0.001; BAI: t=-4.60, p<0.001). Note that this decrease is irrespective of counterbalancing the intervention sequence.

### 4.2. Overall EEG power spectral differences

Power spectral values were compared between first and last days of the 1-week intervention across all 29 electrodes. Both the groups showed significant power changes between the interventions across various frequencies (Fig 4 and 5). SM intervention showed greater increase in power than when the same participants were exposed to silence intervention, with the most prominent difference (based on tfce scores) seen in the gamma band (30-70Hz) (Fig 6). In contrast, VM intervention showed prominent decrease in power across all frequency bands than when the same participants were exposed to silence intervention (Fig 7). The significant change in both the music groups was most evident in the bilateral temporal, parietal and occipital regions. Note that the silence intervention in both the groups was associated with a trend towards increase in all frequency bands.

**Figure 2:**
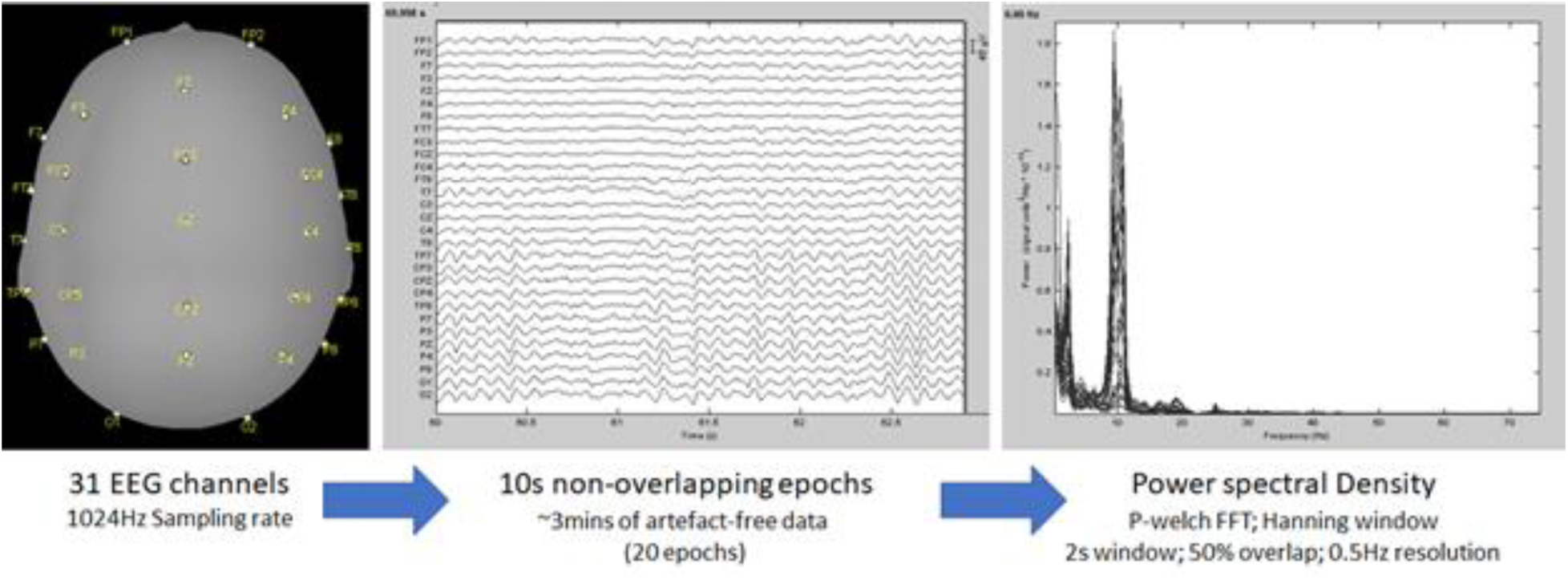
Power spectral analysis flow chart

**Figure 3:**
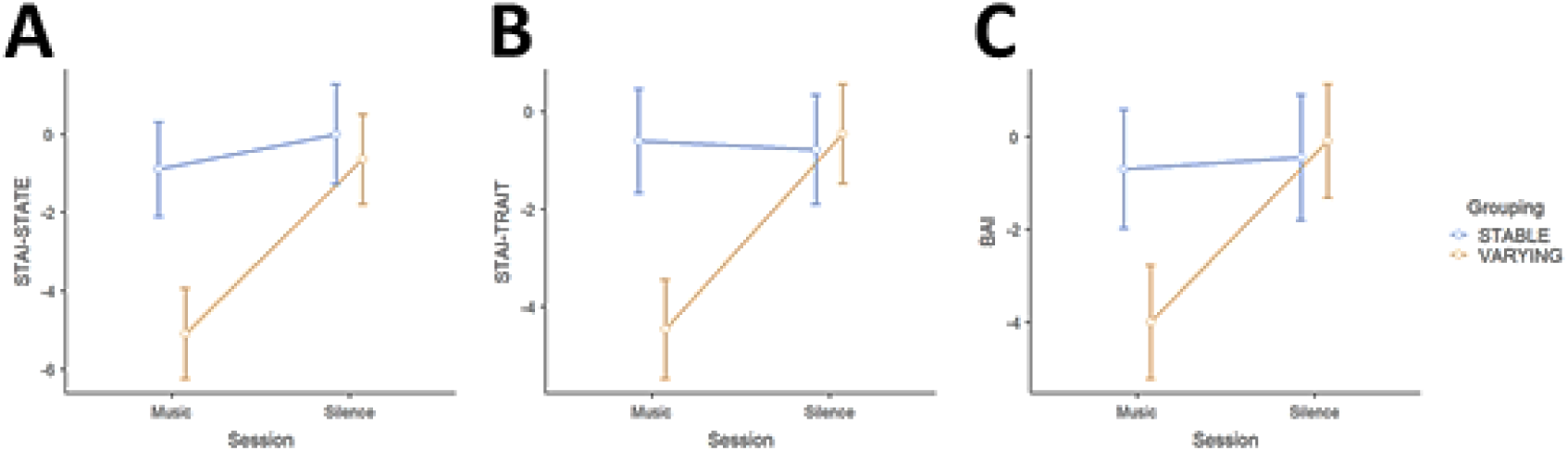
Estimated marginal means and 95% confidence intervals of anxiety scores between the interventions and groups. The scores are based on: (A) State-subscale of State trait anxiety inventory (STAI), (B) Trait-subscale of STAI, and (C) Beck anxiety inventory (BAI). Note that music intervention in Varying music group show significantly lower anxiety scores in all the three measures, irrespective of counterbalancing the intervention sequence.

**Figure 4:**
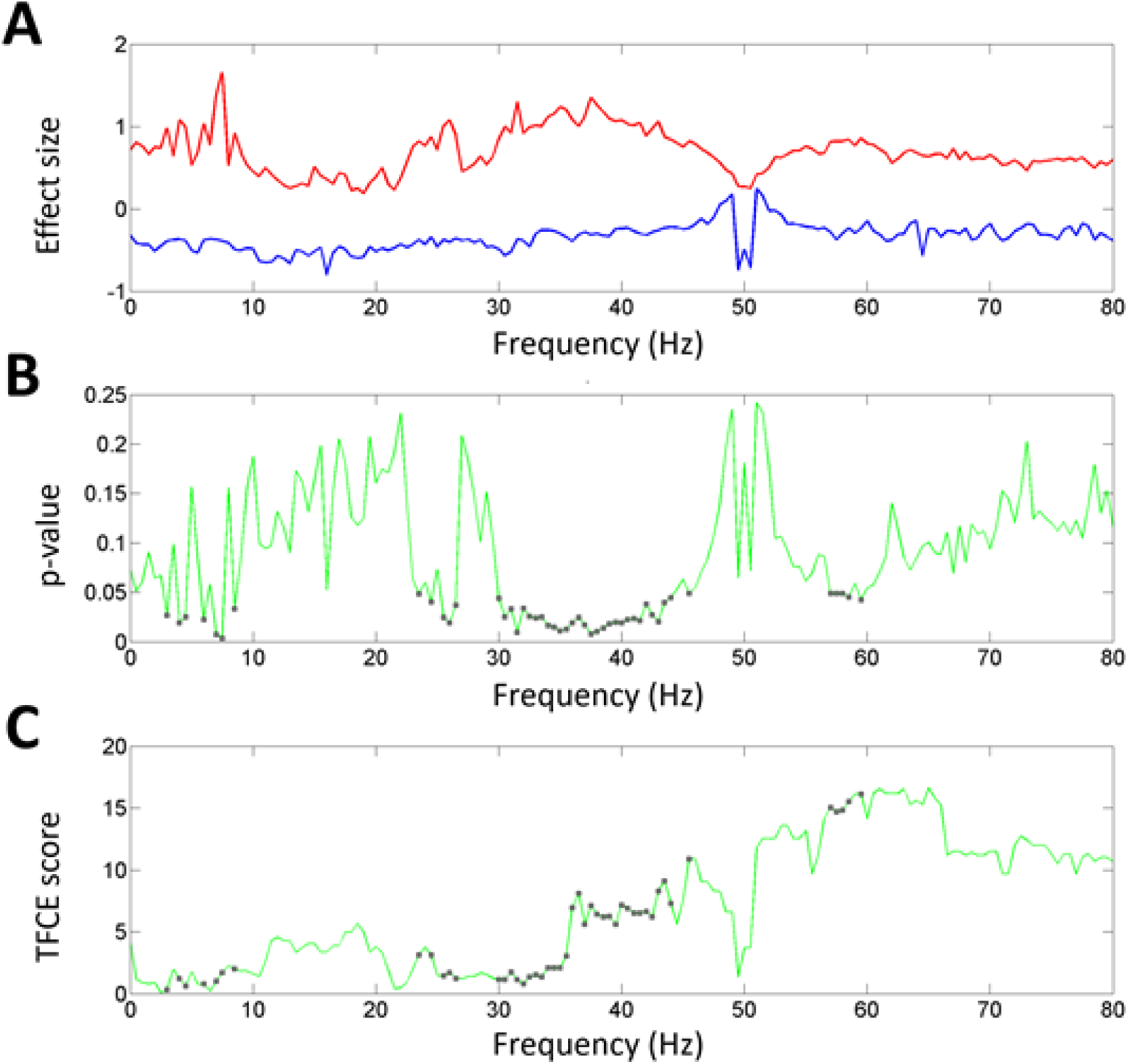
Statistical comparison of power spectral changes between the interventions among Stable music group. Line graphs show the distribution of: (A) effect size of all channels (upper envelope - red; lower envelope - blue), (B) p-value based on cluster statistics, and (C) threshold-free cluster enhancement (tfce) score. The statistics is based on hierarchical general linear model analysis between music and silence interventions. Grey dots on the line graphs correspond to values that are statistically significant at p < 0.05. Note that the significant changes are mostly in higher frequencies.

**Figure 5:**
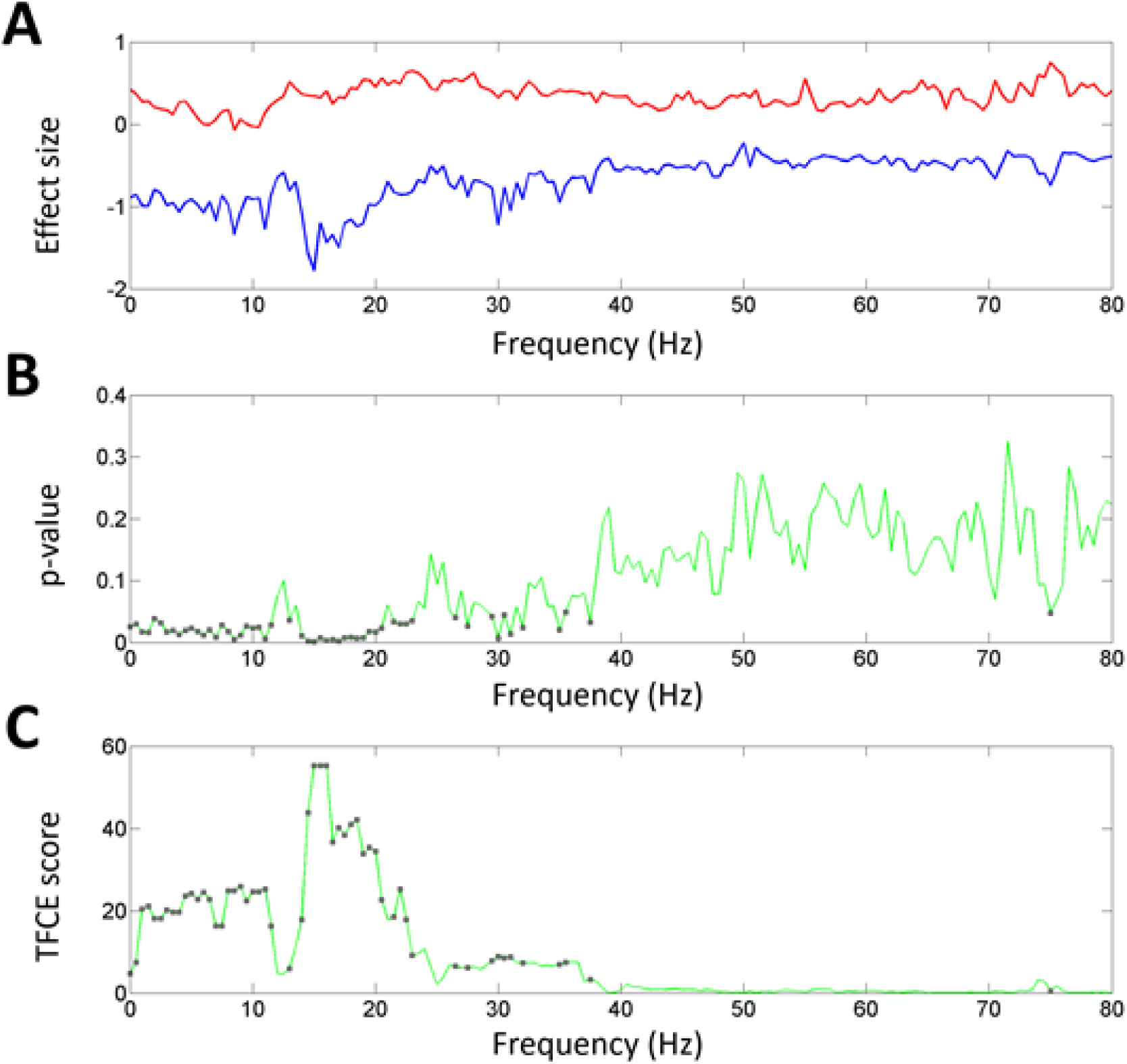
Statistical comparison of power spectral changes between the interventions among Varying music group. Line graphs show the distribution of: (A) effect size of all channels (upper envelope - red; lower envelope - blue), (B) p-value based on cluster statistics, and (C) threshold-free cluster enhancement (tfce) score. The statistics is based on hierarchical general linear model analysis between music and silence interventions. Grey dots on the line graphs correspond to values that are statistically significant at p < 0.05. Note that the significant changes are mostly in lower frequencies.

**Figure 6:**
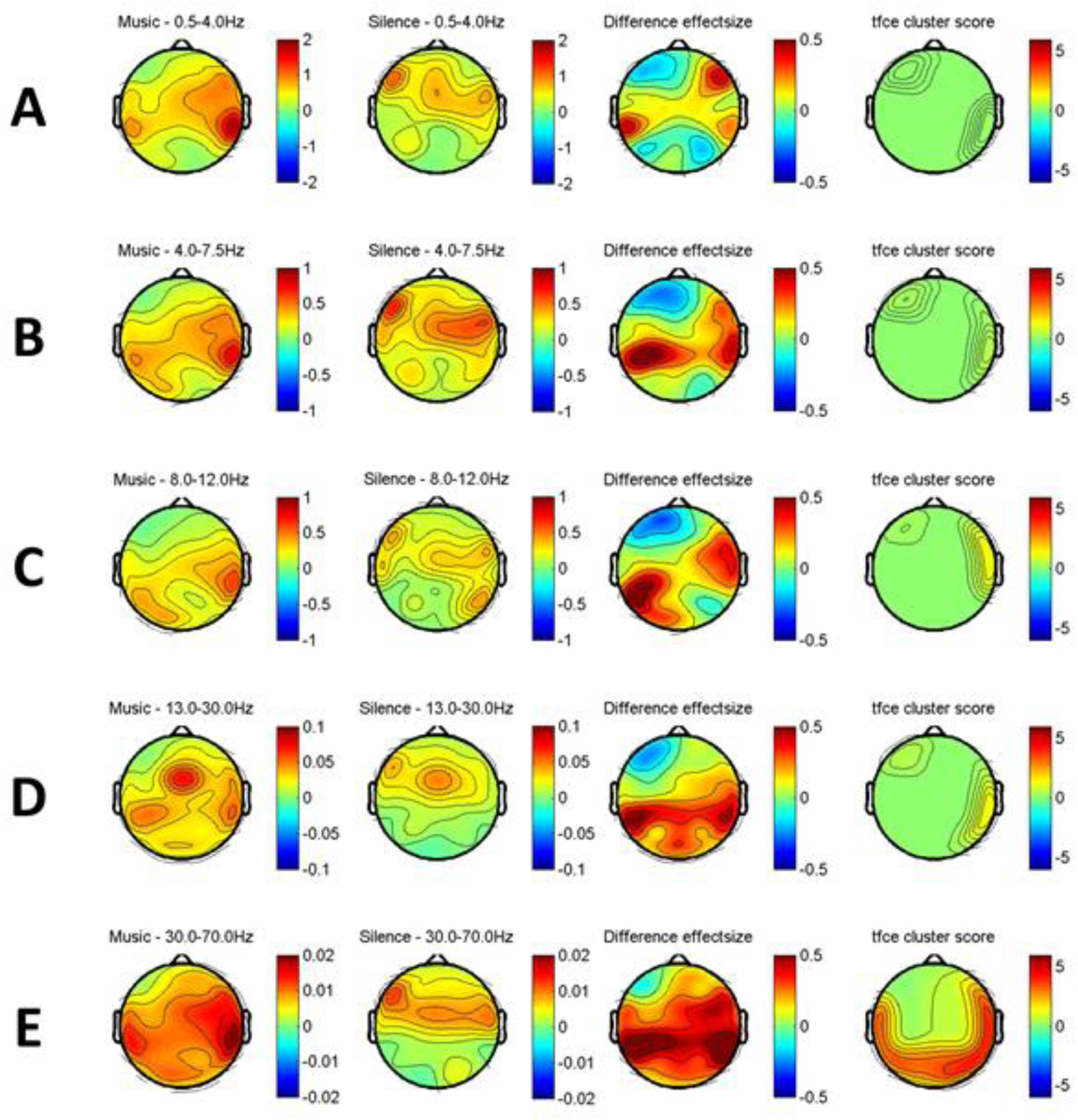
Scalp distribution of statistically significant power spectral changes across the interventions among Stable music group in (A) delta (0.5-4Hz), (B) theta (4-7.5Hz), (C) alpha (8-12Hz), (D) beta (13-30Hz) and (E) gamma (30-80Hz) bands. The scalp maps are generated by averaging the frequency points that showed significant difference in the hierarchical general linear model analysis.The first two columns (from the left) show the average power changes (first-level beta values) associated with music and silence interventions, respectively. The next two columns show the difference in power change between the two interventions in terms of effect size and threshold-free cluster enhancement (tfce) score, respectively (second-level results).

**Figure 7:**
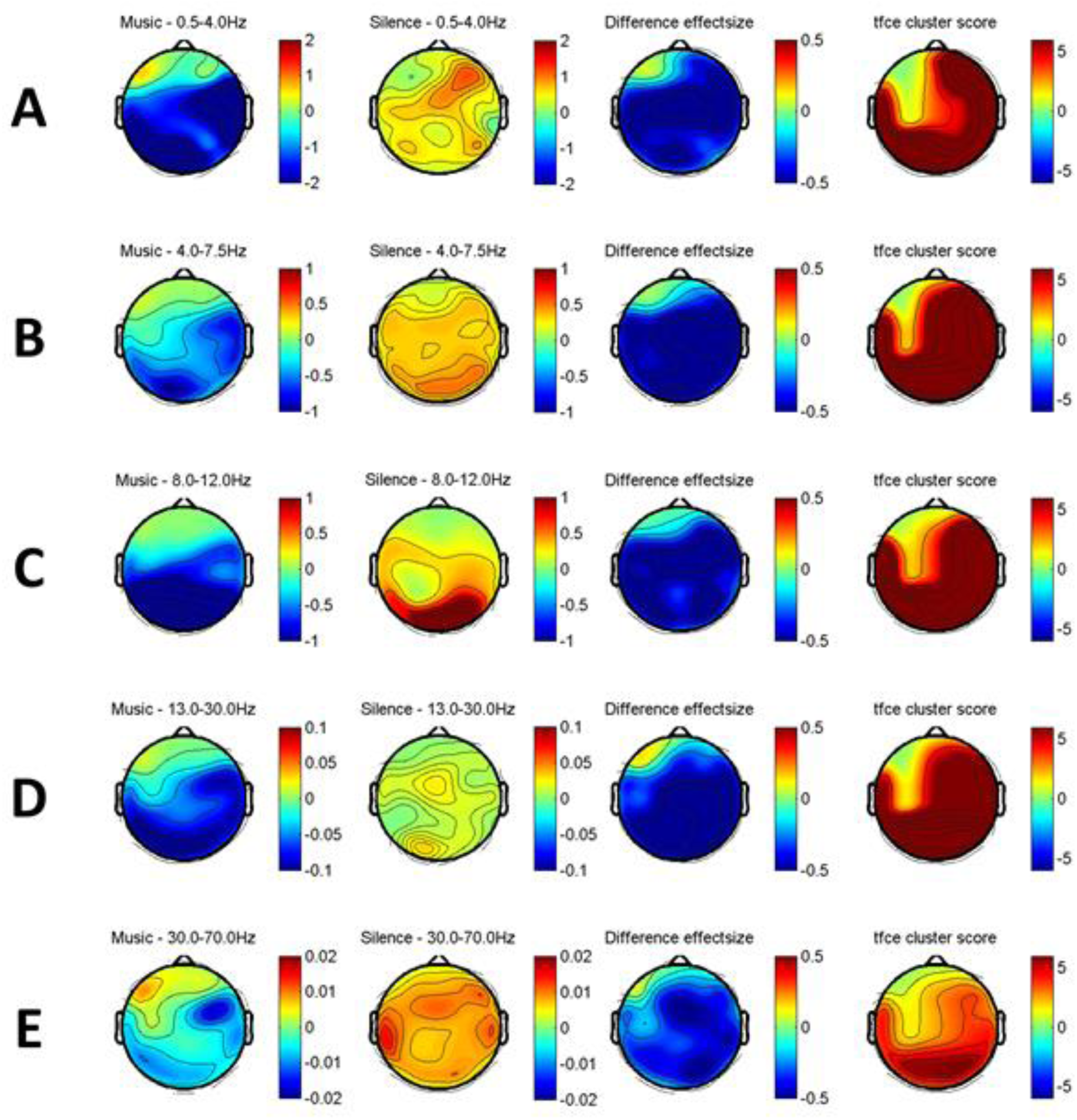
Scalp distribution of statistically significant power spectral changes across the interventions among Varying music group in (A) delta (0.5-4Hz), (B) theta (4-7.5Hz), (C) alpha (8-12Hz), (D) beta (13-30Hz) and (E) gamma (30-80Hz) bands. The scalp maps are generated by averaging the frequency points that showed significant difference in the hierarchical general linear model analysis. The first two columns (from the left) show the average power changes (first-level beta values) associated with music and silence interventions, respectively. The next two columns show the difference in power change between the two interventions in terms of effect size and threshold-free cluster enhancement (tfce) score, respectively (second-level results).

### 4.2. Specific Frontal EEG changes

Frontal asymmetry change was derived from alpha and beta bands combined, to check for frontal hemispheric asymmetry pattern known to be associated with stress and anxiety. Whereas, midline power value (fronto-parietal) was derived from the delta, theta, alpha, beta and gamma bands combined, to investigate the activity of default mode network (DMN) known to be associated with mind wandering. In both groups, more participants showed significant left-sided frontal asymmetry (possibly right prefrontal dominance) and significant reduction in midline power (possibly lower DMN activity) following music intervention, compared to silence intervention (Fig 8A & 8B). Between the groups, more participants in SM group (80% vs 36%) showed left-sided frontal asymmetry whereas more participants in VM group (67% vs 50%) showed midline power decline, when compared to silence (Fig 8C).

**Figure 8:**
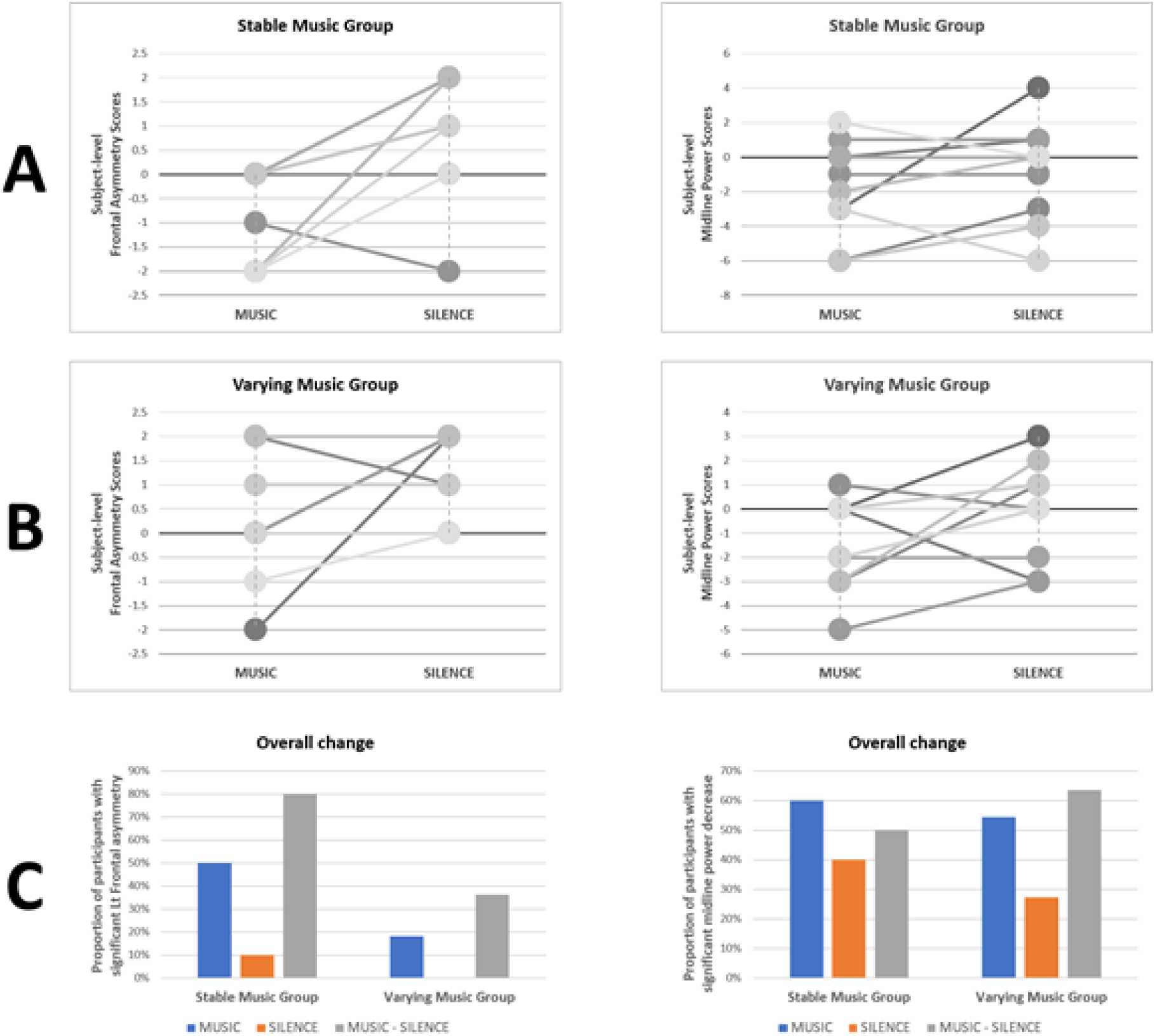
Change in frontal asymmetry (Left column) and midline spectra power (Right column) across the interventions among both groups. Frontal asymmetry change is derived from alpha (8-12Hz) and beta (13-30Hz) bands combined, whereas midline power value (fronto-parietal) is derived from delta (0.5-4Hz), theta (4-7.5Hz), alpha, beta and gamma (30-80Hz) bands combined. (A) and (B) show the significant subject-level changes following music and silence interventions in both groups. Positive values represent significant right-sided asymmetry (Left column) or significant increase in midline power (Right column) and vice versa. (C) shows the proportion of participants with significant left frontal asymmetry and midline power decrease, independently for music and silence interventions as well as for their paired difference (i.e., whether each person showed decrease in midline power during music compared to silence intervention). Within-subject statistical significance between interventions was determined using bootstrap statistics (1000 iterations; 2-tailed significance set at p-value < 0.05).

### 4.3. Correlation between Anxiety scores and Frontal EEG changes

Frontal asymmetry in alpha-beta power showed significant negative correlation between at least one of the anxiety scores, in both the interventions. For music intervention across both groups Frontal asymmetry correlated with STAI-trait (Kendall’s Tau B = -0.45; p = 0.014; Fig 9A) and for silence intervention Frontal asymmetry correlated with BAI (Kendall’s Tau B = -0.41; p = 0.042; Fig 9B).

**Figure 9:**
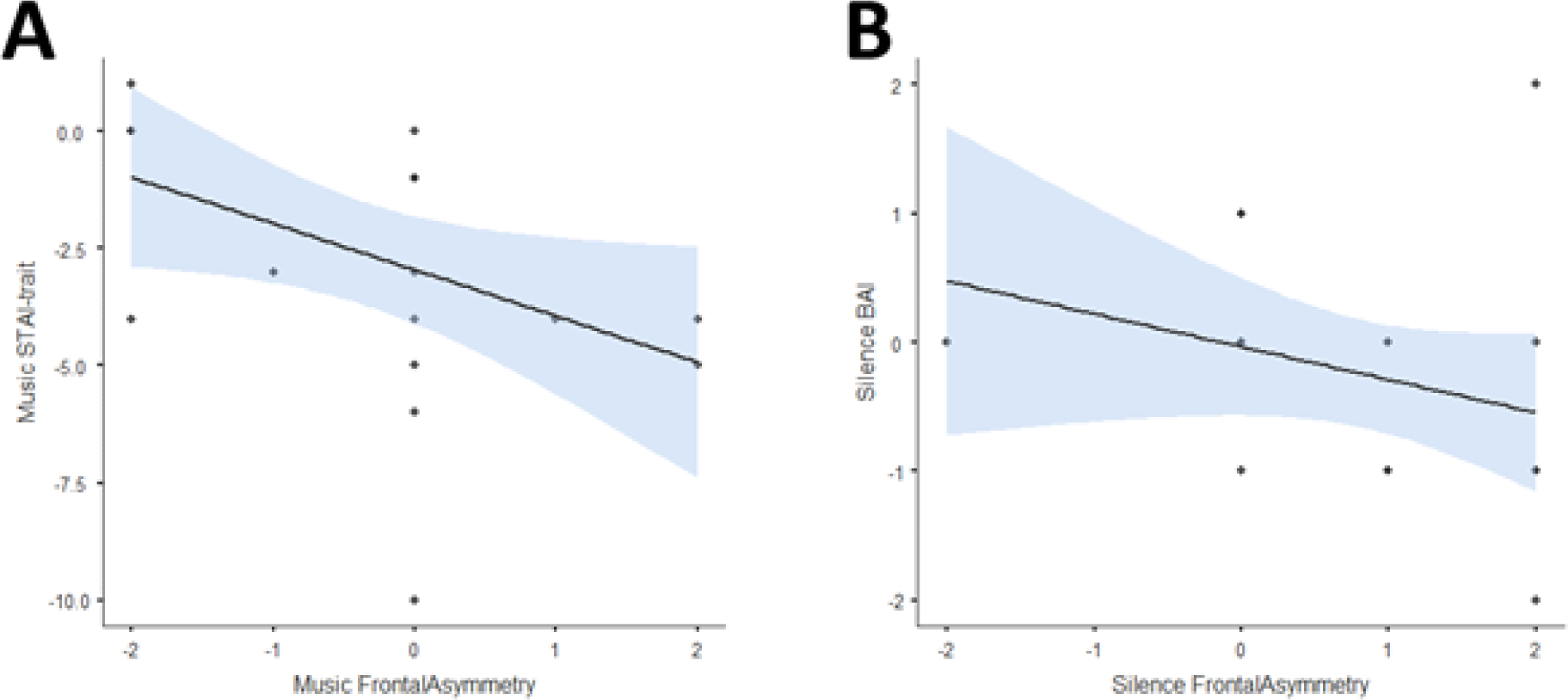
Correlation between frontal asymmetry and anxiety scores that were statistically significant, based on intervention: (A) Music (B) Silence. STAI - State trait anxiety inventory; BAI - Beck anxiety inventory.

### 4.4. Heart rate variability changes

Resting state heart rate variability was compared between first and last day of the 1-week intervention. When silence was used as intervention, significant decrease was observed in relative RR intervals (rrHRV) for both the groups (Table 1 & 2). This change was absent in both the groups, when the music was used as intervention (Table 3 & 4). Other HRV parameters did not show any statistically significant difference.

**Table 1:** Heart rate variability (HRV) changes following silence intervention among stable music group.

|  | Early | Later | Statistics |
| --- | --- | --- | --- |
| rrHRV median | 6.726 ± 2.149 | 5.338 ± 1.972 | Student's t test:<br>$t = 1.955$; $df = 9$; |
|  |  |  | p = 0.082;<br>Cohen's d = 0.618 |
| **rrHRV interquartile range** | 4.330 ± 1.220 | 3.237 ± 1.163 | Student's t test:<br>t = 2.493; df = 9;<br>p = 0.034;<br>Cohen's d = 0.788 |
| **Heart rate (median; bpm)** | 81.314 ± 9.080 | 80.577 ± 11.714 | Student's t test:<br>t = 0.187; df = 9;<br>p = 0.856;<br>Cohen's d = 0.059 |
| **LF/HF ratio** | 0.833 ± 0.323 | 1.030 ± 0.579 | Student's t test:<br>t = -0.953; df = 9;<br>p = 0.365;<br>Cohen's d = -0.301 |

**Table 2:** Heart rate variability (HRV) changes following silence intervention among varying music group.

|  | **Early** | **Later** | **Statistics** |
| --- | --- | --- | --- |
| **rrHRV median** | 7.824 ± 3.791 | 6.188 ± 3.239 | Wilcoxon test:<br>W = 47.00; df = 9;<br>p = 0.049;<br>Cohen's d = 0.513 |
| **rrHRV interquartile range** | 5.435 ± 2.534 | 4.467 ± 1.783 | Student's t test:<br>t = 1.422; df = 9;<br>p = 0.189;<br>Cohen's d = 0.450 |
| **Heart rate (median; bpm)** | 73.108 ± 9.586 | 74.945 ± 10.168 | Student's t test:<br>t = -0.655; df = 9;<br>p = 0.529;<br>Cohen's d = -0.207 |
| **LF/HF ratio** | 0.777 ± 0.371 | 0.860 ± 0.281 | Wilcoxon test:<br>W = 9.00; df = 9;<br>p = 0.064;<br>Cohen's d = -0.430 |

**Table 3:** **Heart rate variability (HRV) changes following music intervention among stable music group**

|  | **Early** | **Later** | **Statistics** |
| --- | --- | --- | --- |
| **rrHRV** | 7.235 ± 3.505 | 6.961 ± 2.952 | Student's t test: |
| **median** |  |  | t = 0.175; df = 9;<br>p = 0.865;<br>Cohen's d = 0.056 |
| **rrHRV**<br>**interquartile range** | 4.612 ± 1.860 | 4.329 ± 2.418 | Student's t test:<br>t = 0.307; df = 9;<br>p = 0.766;<br>Cohen's d = 0.097 |
| **Heart rate (median;**<br>**bpm)** | 80.405 ± 8.380 | 78.279 ± 9.317 | Student's t test:<br>t = 0.621; df = 9;<br>p = 0.550;<br>Cohen's d = 0.196 |
| **LF/HF ratio** | 0.798 ± 0.303 | 0.749 ± 0.274 | Student's t test:<br>t = 0.439; df = 9;<br>p = 0.671;<br>Cohen's d = 0.139 |

**Table 4:** Heart rate variability (HRV) changes following music intervention among varying music group.

|  | **Early** | **Later** | **Statistics** |
| --- | --- | --- | --- |
| **rrHRV**<br>**median** | 6.073 ± 2.216 | 5.973 ± 2.867 | Student's t test:<br>t = 0.144; df = 9;<br>p = 0.889;<br>Cohen's d = 0.046 |
| **rrHRV**<br>**interquartile range** | 4.774 ± 2.460 | 4.812 ± 2.600 | Student's t test:<br>t = -0.056; df = 9;<br>p = 0.956;<br>Cohen's d = -0.018 |
| **Heart rate (median;**<br>**bpm)** | 70.092 ± 9.158 | 73.771 ± 11.727 | Student's t test:<br>t = -1.359; df = 9;<br>p = 0.207;<br>Cohen's d = -0.430 |
| **LF/HF ratio** | 0.971 ± 0.351 | 0.887 ± 0.431 | Wilcoxon test:<br>W = 39.0; df = 9;<br>p = 0.275;<br>Cohen's d = 0.266 |

## Discussion

The present study examined whether adding specific variations in tempo and octave to music (such as a crescendo pattern) can have non-trivial influences on the physiological and psychological mechanisms that are beneficial to stress management. Our results show that participants when exposed to VM showed significant reduction in anxiety, in contrast to SM or silence intervention. A global examination of power spectral changes showed a stark contrast between SM and VM intervention in comparison to silence; former showing greater increase in higher frequencies whereas latter showing prominent decrease especially in lower frequencies, both in bilateral temporo-parieto-occipital regions. A more detailed spectral analysis in frontal region revealed that more participants showed significant left-dominant alpha/beta asymmetry (i.e., greater right brain activation) and decrease in overall midline power (i.e., lower DMN activity) during music intervention, when compared to silence intervention. Interestingly, left asymmetry was more among SM group, whereas, midline power reduction was more among VM group. Both music intervention also didn’t show the reduction in HRV parameters that was associated with silence intervention.

To the best of our knowledge, there are no prior reports on anxiety reduction following a combined manipulation of tempo and octave in a music clip. However, there are many isolated findings that support our observation. For instance, isolated increase in musical tempo is known to stimulate arousal level and vigilance of an individual (C. Hasegawa & Oguri, 2006). With regard to the combinations of tempo and valence of music, a study reported that faster-positive valence music reduce mind wandering (Taruffi et al., 2017), whereas another study reported that slow-positive valence music played during breaks improved the performance in sustained attention tasks (Baldwin & Lewis, 2017). Another study (Mathur, Vijayakumar, Chakrabarti, & Singh, 2015) conducted to assess the emotional response to various *Hindustani Raaga*, reported that the transition from slower-arrhythmic portions (*Alaap*) to faster-rhythmic portions (*Gat*) was associated with change in emotional response; viz., from ‘calm/soothing’ to ‘happy’, respectively. Furthermore, this study found that, irrespective of such influence by tempo/rhythm, all ragas universally generated a ‘calming effect’. Hence, it can be said that although music of a particular *Raaga* is capable of inducing an overall calming effect, its low-level music properties govern the more subtle responses like arousal, sustained attention, emotional response, mind wandering and mindfulness.

We found a global change in power spectral changes (i.e., not limited to a frequency band or scalp location) between music intervention and silence. Although few earlier studies have reported an increase in alpha power over posterior or frontal regions, there has been inconsistency in prior literature in this regard. Increase in power across frequency bands over widespread scalp locations has been reported earlier and was interpreted as a sign of increased internal attention or mind wandering (Jäncke, Kühnis, Rogenmoser, & Elmer, 2015). This is in line with our findings in SM group. So, listening to SM for a week may be causing more mind wandering. Conversely, another study (Lin et al., 2014) had reported a decrease in theta, alpha and beta power, and increase in sympathetic tone after listening to a single session of ‘Mozart music’. The authors of the study reported that their findings suggest an increase in sustained attention and cognitive functions associated with music listening. This supports our findings in VM group. Thus, listening to VM for a week may be causing less mind wandering and more cognitive processing, despite being engrossed into the music. The frequency non-specific nature of our findings could be explained based on prior biophysical studies, which state that each neuron is capable of oscillating at multiple frequencies and that no brain region (in general) display pure brain oscillations (Buzsáki & Draguhn, 2004; Llinás, 1988). Hence, a collective influence on delta, theta, alpha, beta and gamma rhythms could be ascribed to network-level changes in cortico-thalamo systems (Steriade, 2006). To further examine this differential effect on spectral power with both types of music, we focussed on two electrophysiological measures in frontal regions (hemispheric asymmetry and midline power) known to be associated with anxiety/negative mentation and mind wandering, respectively.

Indeed, we found that more subjects who listened to SM than VM (both higher than silence), showed left frontal asymmetry (implying right frontal activation). Several studies in the past using various kinds of music stimuli have explored the hemispheric specialization during music listening. For instance, one study looking at frontal alpha power (Schmidt & Trainor, 2001) reported greater left and right activations while listening to ‘pleasant’ and ‘unpleasant’ music respectively, in a group of undergraduate students. Whereas, another study looking at negative dc-shift fronto-temporal EEG (Altenmüller, Schürmann, Lim, & Parlitz, 2002) presented four different genres of music and reported predominant left hemisphere activation for positive valence stimuli and a more bilateral activation for negative valence stimuli. Physiologically, it has been suggested that lateralization is due to differences in the inputs from the autonomic nervous system (Craig, 2005); right hemisphere predominantly receiving sympathetic inputs associated with greater arousal and negative effects, while the left hemisphere predominantly receiving parasympathetic inputs associated with lesser arousal and positive effects. Thus, right frontal suppression of alpha band power (implying greater right frontal activity) is seen as an index of increased physiological activation and associated with heightened internal attentional states (Jann et al., 2009; Klimesch, 1999). Frontal asymmetry has also been reported as trait effect, with several studies reporting relative increase in right frontal activity in both depression and anxiety disorders (Bruder et al., 1997; Kemp et al., 2010; Mathersul, Williams, Hopkinson, & Kemp, 2008). Heller et al (Heller, Nitschke, Etienne, & Miller, 1997) argues that this pattern seen in anxiety disorders enhances with heightened anxiety arousal. It may also be noted that increased frontal alpha with asymmetry has been linked to creative thinking (Jauk, Benedek, & Neubauer, 2012) and greater right brain activity in resting state is associated with insightful problem solving (Jung-Beeman et al., 2004; Kounios et al., 2008). From the above observations it is pertinent that right brain dominance (reflected by relative left alpha/beta increase) could be a trait sign of persistent negative valence thoughts and anxious arousal, but at the same time can help in creative/insightful ideation.

Another important finding in frontal region was that more subjects who listened to VM than SM (both more that silence), showed significant reduction in midline frontal power, suggesting a greater reduction in DMN activity. Midline structures including posterior cingulate cortex, adjacent precuneus, medical prefrontal cortex, mesial and inferior temporal lobes and inferior parietal lobes represent DMN activity (Broyd et al., 2009; Greicius, Supekar, Menon, & Dougherty, 2009). DMN refers to the constellation of brain regions functionally defined by decreased activation by goal oriented or attention demanding tasks (Buckner, Andrews-Hanna, & Schacter, 2008), which is known to mediate self-referential processing (Whitfield-Gabrieli et al., 2011); viz., remembering one’s past, planning for the future and forming one’s beliefs (Marcus E. Raichle & Snyder, 2007). DMN activity is also attributed to processes ranging from attentional lapses to anxiety to clinical disorders, such as attention-deficit hyperactivity disorder (ADHD) and Alzheimer’s Disease (Buckner et al., 2008; Castellanos et al., 2008; Weissman, Roberts, Visscher, & Woldorff, 2006). Relevant to the current study, DMN activity is positively correlated with mind wandering (Andrews-Hanna, Reidler, Sepulcre, Poulin, & Buckner, 2010; Mason et al., 2007; M. E. Raichle et al., 2001; Simpson, Drevets, Snyder, Gusnard, & Raichle, 2001). Several studies in the past have found significant relations between EEG power changes, DMN activity and mind wandering. Scheeringa et al (Scheeringa et al., 2008) reported a negative correlation between frontal midline theta power and DMN activity. Whereas, Berkovich-Ohana et al (Berkovich-Ohana, Glicksohn, & Goldstein, 2012) showed a reduction in midline gamma power across all midline regions, and Braboszcz et al (Braboszcz & Delorme, 2011) observed reduced theta power over parietal midline area and delta power over frontal midline area during transition between mind wandering to the on-going task. Considering the discrepancies existing across various EEG spectral frequency bands representing DMN activity and mind wandering, Kawashima et al (Kawashima & Kumano, 2017) suggest that EEG frequency that represent mind wandering state may span across various frequency bands and any mid-line power increase irrespective of the frequency can be related to DMN activity and mind wandering. Our findings follow this latter concept and therefore would suggest that listening to VM was associated with lower DMN activation and therefore lesser mind wandering.

An integrated view of our EEG findings could be made based on a recent fMRI-based meta-analysis study that provides a strong empirical model for anxiety, stating that there is significant hypo-connectivity between affective network, executive control network and DMN (Xu et al., 2019). If right dominant activity may be related to affective network and midline power to DMN, then overall decrease in power following music intervention may be related to their effective interaction with executive/cognitive processing, which was observed only in subjects after exposure to VM. So, the poor control of DMN in subjects with higher trait anxiety and its improvement following music intervention was well captured by the EEG measures used in the study.

We did not see any significant difference in the HRV parameters before and after both the music interventions. But, when the same participants underwent silence intervention for the same period, their relative-RR based HRV parameters showed significant difference. Relative-RR based HRV parameters are a robust measure of parasympathetic influence on HRV, and reduction in such resting state vagally-mediated HRV has been associated with poor emotion regulation (Koval et al., 2013; Steinfurth et al., 2018) and poor cognitive flexibility (Colzato, Jongkees, de Wit, van der Molen, & Steenbergen, 2018). Thus, absence of reduction of HRV may be a beneficial effect of music intervention. However, we did not find any significant difference between VM and SM, as seen in our EEG parameters. It is possible that HRV parameters may be measuring physiological markers unrelated to the ones captured by EEG and anxiety scores.

We also have a speculation on how VM and SM differentially modulates the brain activity, from a phenomenology perspective. As music perception requires close monitoring of its temporal evolution, it would force an individual to take a present-centric perspective to engage with it.

Musical portions with high arousals and valence (like the increasing tempo and octave seen in national anthems) can guide a person more into such state and reduce mind wandering. Hasegawa et al (C. Hasegawa & Oguri, 2006) suggest that slow paced music is usually associated with low levels of alertness and vigilance, whereas high tempo music produces a heightened state of vigilance and alertness. Also, Taruffi et al (Taruffi et al., 2017) suggest that DMN activity can be modulated as a function of happy or sad music, quote happy music to be fast paced and sad music to be slow paced. While listening to VM, the slower portions of the music evokes a low vigilance and low alert state that would guide the subjects into deep internal attention (or more mind wandering state), and thereby leading to increased DMN activity. As the tempo and octave increases progressively, the subject’s level of vigilance and alertness increases such that the attention shifts to music and its evoked pleasant feel (a more mindful state), and in the process reducing the DMN activity. As such incremental variations in the tempo and octave repeats multiple times in the VM clip, the subjects are made to effortlessly switch between mind wandering and mindful states, reducing the occurrence of negative ruminating thoughts that are often associated with longer periods of mind wandering. This detachment from the irrelevant stimuli perhaps is also one of the reasons for relaxation alongside the soothing effects of music. On the contrary, we argue that study subjects perceive SM to be unpleasant and sad, enter heightened internal attentional states that may be seen as increased activation of right frontal regions. Because of the absence of subtle novelties in the music clip, they tend to maintain the same train of thought, fail to breakthrough and direct the attention to stimulus thus making them vulnerable to maladaptive ruminating thoughts. This extensive mind wandering will be associated with increased DMN activation, and when combined with ruminating thoughts suppresses one’s creative thinking, decreases vigilance and alertness and decreases overall task performance. Hence, we do not see any significant post-treatment reduction of scores in the psychological assessments in SM group.

Our study has some limitations. The sample size is only 10-11 per group. But a cross-over design ensured that the same subjects could be used as their own controls, and compensate for the lower sample. However, future studies need to replicate our findings in other populations and other genre of music. We did not choose music based on our subject’s taste, which could have produced stronger effects. But having a common music custom-composed for the study makes it possible to avoid the confounding effects due stimulus variations and strengthens the control over experimental intervention, without compromising on the naturality of musical clip.

The current study looked only into two levels of variations in tempo and octave (incremental and non-incremental), but it would be interesting to explore the effects of other common music variations. We hope to explore the same in subsequent studies.

Thus, we conclude that music without specific variations in musical properties such as tempo and octave can cause extensive mind wandering, increased ruminating thoughts, and when practiced over long periods may have detrimental consequences on the mental health of an individual. Whereas, the incremental variations in tempo and octave induces a controlled switching between “attention to self” during slower portions and “attention to music” during faster portions of music. This ‘controlled Mind wandering’ state can help train one’s creative ideation concurrently maintaining required attentional focus to successfully accomplish the task in hand. Maintaining high levels of vigilance and alertness can be very demanding and stressful and medical profession demands a sound attentional focus on the task and extensive mind wandering may result in errors in medical practice. Given the fact that mind wandering is difficult to control in an ever-stressful environment it is indispensable to introduce techniques that promote mindfulness to avoid fatigue, low mood and various mental health disorders within the medical community.

Therefore, our study provides additional scientific support for introducing Music therapy in standard medical education curriculum to reduce the burden of anxiety related disorders and proposes the importance of choosing appropriate music clips for easy yet effective control of mind wandering.

## Acknowledgement

We thank Dr Ganne Chaithanya (Department of Neurology, Thomas Jefferson University, Philadelphia, United States) and Dr N R Ramesh Masthi (Department of Community Medicine, KIMS Bengaluru) for their valuable inputs during the inception of the study. We also thank the musicians who prepared the music clip used in this study: TKV Ramanujacharyulu (Violin) and Trichy Harikumar (Mridangam).

## REFERENCES

Altenmüller, E., Schürmann, K., Lim, V. K., & Parlitz, D. (2002). Hits to the left, flops to the right: different emotions during listening to music are reflected in cortical lateralisation patterns. Neuropsychologia, 40(13), 2242–2256.

Amezcua, C., Guevara, M. A., & Ramos-Loyo, J. (2005). Effects of Musical Tempi on Visual Attention Erps. International Journal of Neuroscience, 115(2), 193–206. https://doi.org/10.1080/00207450590519094

Andrews-Hanna, J. R., Reidler, J. S., Sepulcre, J., Poulin, R., & Buckner, R. L. (2010). Functional-Anatomic Fractionation of the Brain’s Default Network. Neuron, 65(4), 550–562. https://doi.org/10.1016/j.neuron.2010.02.005

Anuradha, R., Dutta, R., Raja, J. D., Sivaprakasam, P., & Patil, A. B. (2017). Stress and Stressors among Medical Undergraduate Students: A Cross-sectional Study in a Private Medical College in Tamil Nadu. Indian Journal of Community Medicine: Official Publication of Indian Association of Preventive & Social Medicine, 42(4), 222–225. https://doi.org/10.4103/ijcm.IJCM_287_16

Baldwin, C. L., & Lewis, B. A. (2017). Positive valence music restores executive control over sustained attention. PLOS ONE, 12(11), e0186231. https://doi.org/10.1371/journal.pone.0186231

Beck, A. T., Epstein, N., Brown, G., & Steer, R. A. (1988). An inventory for measuring clinical anxiety: psychometric properties. Journal of Consulting and Clinical Psychology, 56(6), 893–897.

Berkovich-Ohana, A., Glicksohn, J., & Goldstein, A. (2012). Mindfulness-induced changes in gamma band activity – Implications for the default mode network, self-reference and attention. Clinical Neurophysiology, 123(4), 700–710. https://doi.org/10.1016/j.clinph.2011.07.048

Braboszcz, C., & Delorme, A. (2011). Lost in thoughts: Neural markers of low alertness during mind wandering. NeuroImage, 54(4), 3040–3047. https://doi.org/10.1016/j.neuroimage.2010.10.008

Broyd, S. J., Demanuele, C., Debener, S., Helps, S. K., James, C. J., & Sonuga-Barke, E. J. (2009). Default-mode brain dysfunction in mental disorders: a systematic review. Neuroscience & Biobehavioral Reviews, 33(3), 279–296.

Bruder, G. E., Fong, R., Tenke, C. E., Leite, P., Towey, J. P., Stewart, J. E., … Quitkin, F. M. (1997). Regional brain asymmetries in major depression with or without an anxiety disorder: a quantitative electroencephalographic study. Biological Psychiatry, 41(9), 939–948.

Buckner, R. L., Andrews-Hanna, J. R., & Schacter, D. L. (2008). The brain’s default network. Annals of the New York Academy of Sciences, 1124(1), 1–38.

Burns, J. L., Labbé, E., Arke, B., Capeless, K., Cooksey, B., Steadman, A., & Gonzales, C. (2002). The effects of different types of music on perceived and physiological measures of stress. Journal of Music Therapy, 39(2), 101–116.

Buzsáki, G., & Draguhn, A. (2004). Neuronal oscillations in cortical networks. Science, 304(5679), 1926–1929.

Castellanos, F. X., Margulies, D. S., Kelly, C., Uddin, L. Q., Ghaffari, M., Kirsch, A., … Milham, M. P. (2008). Cingulate-Precuneus Interactions: A New Locus of Dysfunction in Adult Attention-Deficit/Hyperactivity Disorder. Biological Psychiatry, 63(3), 332–337. https://doi.org/10.1016/j.biopsych.2007.06.025

Cepeda, M. S., Carr, D. B., Lau, J., & Alvarez, H. (2006). Music for pain relief. The Cochrane Database of Systematic Reviews, (2), CD004843. https://doi.org/10.1002/14651858.CD004843.pub2

Colzato, L. S., Jongkees, B. J., de Wit, M., van der Molen, M. J. W., & Steenbergen, L. (2018). Variable heart rate and a flexible mind: Higher resting-state heart rate variability predicts better task-switching. *Cognitive, Affective*, & Behavioral Neuroscience, 18(4), 730–738. https://doi.org/10.3758/s13415-018-0600-x

Craig, A. D. (2005). Forebrain emotional asymmetry: a neuroanatomical basis? Trends in Cognitive Sciences, 9(12), 566–571.

Daly, I., Hallowell, J., Hwang, F., Kirke, A., Malik, A., Roesch, E., … Nasuto, S. J. (2014). Changes in music tempo entrain movement related brain activity. 2014 36th Annual International Conference of the IEEE Engineering in Medicine and Biology Society, 4595–4598. https://doi.org/10.1109/EMBC.2014.6944647

Delorme, A., & Makeig, S. (2004). EEGLAB: an open source toolbox for analysis of single-trial EEG dynamics including independent component analysis. Journal of Neuroscience Methods, 134(1), 9–21. https://doi.org/10.1016/j.jneumeth.2003.10.009

Dileo, C., & Bradt, J. (2009). Music therapy: applications to stress management. In P. M. Lehrer, R. L. Woolfolk, & W. E. Sime (Eds.), Principles and practice of stress management (Third). New York; London: Guilford.

Epstein, R. M. (1999). Mindful practice. JAMA, 282(9), 833–839.

Firth-Cozens, J. (1999). The stresses of medical training. In J. Firth-Cozens & R. Payne (Eds.), Stress in health professionals: psychological and organisational causes and interventions. Chichester ; New York: John Wiley and Sons Ltd.

Fish, C., & Nies, M. A. (1996). Health Promotion Needs of Students in a College Environment. Public Health Nursing, 13(2), 104–111. https://doi.org/10.1111/j.1525-1446.1996.tb00227.x

Greicius, M. D., Supekar, K., Menon, V., & Dougherty, R. F. (2009). Resting-state functional connectivity reflects structural connectivity in the default mode network. Cerebral Cortex, 19(1), 72–78.

Gupta, U., & Gupta, B. S. (2005). Psychophysiological responsivity to Indian instrumental music. Psychology of Music, 33(4), 363–372. https://doi.org/10.1177/0305735605056144

Harvey, A. W. (1987). Utilizing music as a tool for healing. In R. R. Pratt (Ed.), The Fourth International Symposium on Music--Rehabilitation and Human Well-Being: August 1-5, 1985, New York City (pp. 73–87). Lanham: University Press of America.

Hasegawa, C., & Oguri, K. (2006). The effects of specific musical stimuli on driver’s drowsiness. 2006 IEEE Intelligent Transportation Systems Conference, 817–822. https://doi.org/10.1109/ITSC.2006.1706844

Hasegawa, H., Uozumi, T., & Ono, K. (2003). Physiological evaluation of music effect for the mental workload. Proceedings of 6th Asian Design International Conference Tsukuba, Japan. Citeseer.

Heller, W., Nitschke, J. B., Etienne, M. A., & Miller, G. A. (1997). Patterns of regional brain activity differentiate types of anxiety. Journal of Abnormal Psychology, 106(3), 376.

Henry, E. O. (2002). The Rationalization of Intensity in Indian Music. Ethnomusicology, 46(1), 33. https://doi.org/10.2307/852807

Hevner, K. (1937). The Affective Value of Pitch and Tempo in Music. The American Journal of Psychology, 49(4), 621–630.

Iqbal, S., Gupta, S., & Venkatarao, E. (2015). Stress, anxiety and depression among medical undergraduate students and their socio-demographic correlates. The Indian Journal of Medical Research, 141(3), 354–357.

Iwanaga, M., Ikeda, M., & Iwaki, T. (1996). The Effects of Repetitive Exposure to Music on Subjective and Physiological Responses. Journal of Music Therapy, 33(3), 219–230. https://doi.org/10.1093/jmt/33.3.219

Jäncke, L., Kühnis, J., Rogenmoser, L., & Elmer, S. (2015). Time course of EEG oscillations during repeated listening of a well-known aria. Frontiers in Human Neuroscience, 9. https://doi.org/10.3389/fnhum.2015.00401

Jann, K., Dierks, T., Boesch, C., Kottlow, M., Strik, W., & Koenig, T. (2009). BOLD correlates of EEG alpha phase-locking and the fMRI default mode network. Neuroimage, 45(3), 903–916.

Jauk, E., Benedek, M., & Neubauer, A. C. (2012). Tackling creativity at its roots: Evidence for different patterns of EEG alpha activity related to convergent and divergent modes of task processing. International Journal of Psychophysiology, 84(2), 219–225. https://doi.org/10.1016/j.ijpsycho.2012.02.012

Julian, L. J. (2011). Measures of anxiety: State-Trait Anxiety Inventory (STAI), Beck Anxiety Inventory (BAI), and Hospital Anxiety and Depression Scale-Anxiety (HADS-A). Arthritis Care & Research, 63(S11), S467–S472. https://doi.org/10.1002/acr.20561

Jung-Beeman, M., Bowden, E. M., Haberman, J., Frymiare, J. L., Arambel-Liu, S., Greenblatt, R., … Kounios, J. (2004). Neural Activity When People Solve Verbal Problems with Insight. PLoS Biology, 2(4), e97. https://doi.org/10.1371/journal.pbio.0020097

Kawashima, I., & Kumano, H. (2017). Prediction of Mind-Wandering with Electroencephalogram and Non-linear Regression Modeling. Frontiers in Human Neuroscience, 11, 365. https://doi.org/10.3389/fnhum.2017.00365

Kellaris, J. J., & Kent, R. J. (1991). Exploring Tempo and Modality Effects, on Consumer Responses to Music. *ACR North American Advances*, *NA-18*. Retrieved from http://acrwebsite.org/volumes/7168/volumes/v18/NA-18

Kemp, A. H., Griffiths, K., Felmingham, K. L., Shankman, S. A., Drinkenburg, W., Arns, M., … Bryant, R. A. (2010). Disorder specificity despite comorbidity: resting EEG alpha asymmetry in major depressive disorder and post-traumatic stress disorder. Biological Psychology, 85(2), 350–354.

Klimesch, W. (1999). EEG alpha and theta oscillations reflect cognitive and memory performance: a review and analysis. Brain Research Reviews, 29(2–3), 169–195.

Kounios, J., Fleck, J. I., Green, D. L., Payne, L., Stevenson, J. L., Bowden, E. M., & Jung-Beeman, M. (2008). The origins of insight in resting-state brain activity. Neuropsychologia, 46(1), 281–291. https://doi.org/10.1016/j.neuropsychologia.2007.07.013

Koval, P., Ogrinz, B., Kuppens, P., Van den Bergh, O., Tuerlinckx, F., & Sütterlin, S. (2013). Affective Instability in Daily Life Is Predicted by Resting Heart Rate Variability. PLoS ONE, 8(11), e81536. https://doi.org/10.1371/journal.pone.0081536

Kumar, G., Jain, A., & Hegde, S. (2012). Prevalence of depression and its associated factors using Beck Depression Inventory among students of a medical college in Karnataka. Indian Journal of Psychiatry, 54(3), 223. https://doi.org/10.4103/0019-5545.102412

Kumar, V., Talwar, R., & Raut, D. K. (2014). Psychological distress, general self-efficacy and psychosocial adjustments among first year medical college students in New Delhi, India. South East Asia Journal of Public Health, 3(2). https://doi.org/10.3329/seajph.v3i2.20038

Lazarus, R. S., & Folkman, S. (1984). Stress, Appraisal, and Coping. New York: Springer Publishing Company.

Lee, J., & Graham, A. V. (2001). Students’ perception of medical school stress and their evaluation of a wellness elective. Medical Education, 35(7), 652–659.

Lesiuk, T. (2015). The Effect of Mindfulness-Based Music Therapy on Attention and Mood in Women Receiving Adjuvant Chemotherapy for Breast Cancer: A Pilot Study. Oncology Nursing Forum, 42(3), 276–282. https://doi.org/10.1188/15.ONF.276-282

Lin, L.-C., Ouyang, C.-S., Chiang, C.-T., Wu, R.-C., Wu, H.-C., & Yang, R.-C. (2014). Listening to Mozart K.448 decreases electroencephalography oscillatory power associated with an increase in sympathetic tone in adults: a post-intervention study. JRSM Open, 5(10), 205427041455165. https://doi.org/10.1177/2054270414551657

Llinás, R. R. (1988). The intrinsic electrophysiological properties of mammalian neurons: insights into central nervous system function. Science, 242(4886), 1654–1664.

Ma, W., Lai, Y., Yuan, Y., Wu, D., & Yao, D. (2012). Electroencephalogram variations in the α band during tempo-specific perception: NeuroReport, 23(3), 125–128. https://doi.org/10.1097/WNR.0b013e32834e7eac

Mason, M. F., Norton, M. I., Van Horn, J. D., Wegner, D. M., Grafton, S. T., & Macrae, C. N. (2007). Wandering Minds: The Default Network and Stimulus-Independent Thought. Science, 315(5810), 393–395. https://doi.org/10.1126/science.1131295

Mathersul, D., Williams, L. M., Hopkinson, P. J., & Kemp, A. H. (2008). Investigating models of affect: Relationships among EEG alpha asymmetry, depression, and anxiety. Emotion, 8(4), 560.

Mathur, A., Vijayakumar, S. H., Chakrabarti, B., & Singh, N. C. (2015). Emotional responses to Hindustani raga music: the role of musical structure. Frontiers in Psychology, 6. https://doi.org/10.3389/fpsyg.2015.00513

Mullen, T., Kothe, C., Chi, Y. M., Ojeda, A., Kerth, T., Makeig, S., … Tzyy-Ping Jung. (2013). Real-time modeling and 3D visualization of source dynamics and connectivity using wearable EEG. 2013 35th Annual International Conference of the IEEE Engineering in Medicine and Biology Society (EMBC), 2184–2187. https://doi.org/10.1109/EMBC.2013.6609968

Nilsson, U., Unosson, M., & Rawal, N. (2005). Stress reduction and analgesia in patients exposed to calming music postoperatively: a randomized controlled trial. European Journal of Anaesthesiology, 22(2), 96–102.

Oakes, S. (2003). Musical tempo and waiting perceptions. Psychology and Marketing, 20(8), 685–705. https://doi.org/10.1002/mar.10092

Pan, J., & Tompkins, W. J. (1985). A Real-Time QRS Detection Algorithm. *IEEE Transactions on Biomedical Engineering*, BME*-*32(3), 230–236. https://doi.org/10.1109/TBME.1985.325532

Pernet, C. R., Chauveau, N., Gaspar, C., & Rousselet, G. A. (2011). LIMO EEG: A Toolbox for Hierarchical LInear MOdeling of ElectroEncephaloGraphic Data. Computational Intelligence and Neuroscience, 2011, 1–11. https://doi.org/10.1155/2011/831409

Phipps, M. A., Carroll, D. L., & Tsiantoulas, A. (2010). Music as a therapeutic intervention on an inpatient neuroscience unit. Complementary Therapies in Clinical Practice, 16(3), 138–142. https://doi.org/10.1016/j.ctcp.2009.12.001

Raichle, M. E., MacLeod, A. M., Snyder, A. Z., Powers, W. J., Gusnard, D. A., & Shulman, G. L. (2001). A default mode of brain function. Proceedings of the National Academy of Sciences, 98(2), 676–682. https://doi.org/10.1073/pnas.98.2.676

Raichle, Marcus E., & Snyder, A. Z. (2007). A default mode of brain function: A brief history of an evolving idea. NeuroImage, 37(4), 1083–1090. https://doi.org/10.1016/j.neuroimage.2007.02.041

Rigg, M. G. (1964). The Mood Effects of Music: A Comparison of Data from Four Investigators. The Journal of Psychology, 58(2), 427–438. https://doi.org/10.1080/00223980.1964.9916765

Scheeringa, R., Bastiaansen, M. C. M., Petersson, K. M., Oostenveld, R., Norris, D. G., & Hagoort, P. (2008). Frontal theta EEG activity correlates negatively with the default mode network in resting state. International Journal of Psychophysiology, 67(3), 242–251. https://doi.org/10.1016/j.ijpsycho.2007.05.017

Schmidt, L. A., & Trainor, L. J. (2001). Frontal brain electrical activity (EEG) distinguishes valence and intensity of musical emotions. Cognition & Emotion, 15(4), 487–500. https://doi.org/10.1080/02699930126048

Sedghamiz, H. (2014). Matlab Implementation of Pan Tompkins ECG QRS detector. [Data set]. https://doi.org/10.13140/RG.2.2.14202.59841

Selye, H. (1982). History of the stress concept. In L. Goldberger & S. Breznitz (Eds.), Handbook of stress : theoretical and clinical aspects (1st ed, pp. 7–20). New York : Free Press.

Shaw, J. A. (2003). Children exposed to war/terrorism. Clinical Child and Family Psychology Review, 6(4), 237–246.

Simpson, J. R., Drevets, W. C., Snyder, A. Z., Gusnard, D. A., & Raichle, M. E. (2001). Emotion-induced changes in human medial prefrontal cortex: II. During anticipatory anxiety. Proceedings of the National Academy of Sciences, 98(2), 688–693. https://doi.org/10.1073/pnas.98.2.688

Smallwood, J., Fishman, D. J., & Schooler, J. W. (2007). Counting the cost of an absent mind: mind wandering as an underrecognized influence on educational performance. Psychonomic Bulletin & Review, 14(2), 230–236.

Smallwood, J., Fitzgerald, A., Miles, L. K., & Phillips, L. H. (2009). Shifting moods, wandering minds: negative moods lead the mind to wander. *Emotion (Washington*, D.C.), 9(2), 271–276. https://doi.org/10.1037/a0014855

Smallwood, J., Mrazek, M. D., & Schooler, J. W. (2011). Medicine for the wandering mind: mind wandering in medical practice. Medical Education, 45(11), 1072–1080. https://doi.org/10.1111/j.1365-2923.2011.04074.x

Smallwood, J., O’Connor, R. C., Sudbery, M. V., & Obonsawin, M. (2007). Mind-wandering and dysphoria. Cognition & Emotion, 21(4), 816–842. https://doi.org/10.1080/02699930600911531

Smith, S., & Nichols, T. (2009). Threshold-free cluster enhancement: Addressing problems of smoothing, threshold dependence and localisation in cluster inference. NeuroImage, 44(1), 83–98. https://doi.org/10.1016/j.neuroimage.2008.03.061

Spielberger, C. D., Gorsuch, R. L., Lushene, R., Vagg, P. R., & Jacobs, G. A. (1983). Manual for the State-Trait Anxiety Inventory. Palo Alto, CA: Consulting Psychologists Press.

Steinfurth, E. C. K., Wendt, J., Geisler, F., Hamm, A. O., Thayer, J. F., & Koenig, J. (2018). Resting State Vagally-Mediated Heart Rate Variability Is Associated With Neural Activity During Explicit Emotion Regulation. Frontiers in Neuroscience, 12, 794. https://doi.org/10.3389/fnins.2018.00794

Steriade, M. (2006). Grouping of brain rhythms in corticothalamic systems. Neuroscience, 137(4), 1087–1106.

Tadel, F., Baillet, S., Mosher, J. C., Pantazis, D., & Leahy, R. M. (2011). Brainstorm: A User-Friendly Application for MEG/EEG Analysis. Computational Intelligence and Neuroscience, 2011, 1–13. https://doi.org/10.1155/2011/879716

Taruffi, L., Pehrs, C., Skouras, S., & Koelsch, S. (2017). Effects of Sad and Happy Music on Mind-Wandering and the Default Mode Network. Scientific Reports, 7(1). https://doi.org/10.1038/s41598-017-14849-0

Vollmer, M. (2015). A robust, simple and reliable measure of heart rate variability using relative RR intervals. 2015 Computing in Cardiology Conference (CinC), 609–612. https://doi.org/10.1109/CIC.2015.7410984

Weissman, D. H., Roberts, K. C., Visscher, K. M., & Woldorff, M. G. (2006). The neural bases of momentary lapses in attention. Nature Neuroscience, 9(7), 971–978. https://doi.org/10.1038/nn1727

Whitfield-Gabrieli, S., Moran, J. M., Nieto-Castañón, A., Triantafyllou, C., Saxe, R., & Gabrieli, J.D. (2011). Associations and dissociations between default and self-reference networks in the human brain. Neuroimage, 55(1), 225–232.

Xu, J., Van Dam, N. T., Feng, C., Luo, Y., Ai, H., Gu, R., & Xu, P. (2019). Anxious brain networks: A coordinate-based activation likelihood estimation meta-analysis of resting-state functional connectivity studies in anxiety. Neuroscience & Biobehavioral Reviews, 96, 21–30. https://doi.org/10.1016/j.neubiorev.2018.11.005

